# Exosomes for facial repair, regeneration and rejuvenation: comparison between different commercial products

**DOI:** 10.1101/2024.12.07.627351

**Authors:** Antonio Salvaggio, Anna Privitera, Greta Ferruggia, Massimo Zimbone, Maria Violetta Brundo

## Abstract

Exosomes have been studied as a potential therapeutic option for improving skin texture. The mode of action of exosomes for skin texture likely involves several mechanisms, including stimulation of collagen production, promotion of skin cell proliferation; reduction of oxidative stress; regulation of inflammatory and antioxidant responses; extracellular matrix remodeling, which is important for maintaining skin structure and function. Aim of this study is to compare the *in vitro* effects of some commercial products that use new technologies containing plant derived nanovesicles from *Rosa damascena* callus, *Centella asiatica* callus, *Euphorbia supina* stem and exosomes from bovine colostrum passively loaded with growth factors and cytokines purified from bovine colostrum, for facial repair, regeneration and rejuvenation. Of all the products tested, only product containing exosomes purified from colostrum gave very encouraging results in terms of effects on proliferation, cell viability and wound repair. The other products tested gave results comparable to the untreated samples.

## 1. Introduction

In the last decade, interest in extracellular vesicles has grown considerably, in particular since the key role of exosomes in intercellular communication has been demonstrated [1,2]. Exosomes are spherical extracellular nanovesicles with an endosomal origin with sizes ranging from 30 to 100 nm [3], as transporters of different bioactive molecules including proteins and lipids, can take part in different physiological mechanisms by transferring their contents to recipient cells, thus acting as extracellular messengers in cell-to-cell communication, but they are also attracting attention as potential therapeutic tools and positive effects on the regeneration of many tissues [3,4]. In this context, exosomes are attracting growing interest thanks to their relevance in the modulation of cellular processes (physiological and pathological), as well as for their potential beneficial effects on the organism, especially anti-inflammatory, anti-tumor, tissue regeneration and modulation of commensal microbiota. Furthermore, it is hypothesized to also use them as transporters of active molecules, so as to exploit them to improve the absorption and effectiveness of drugs thanks to their low immunogenicity and high stability in the gastrointestinal tract [5]. It is widely accepted that the therapeutic potential of several components purified from bovine colostrum may be mediated largely by paracrine factors, and exosomes might be the main components of such paracrine factors [6]. Thus, exosomes derived from bovine colostrum, as they induce the repair of damaged tissues, represent a relevant therapeutic option in regenerative medicine and have a more promising future [7-11], especially when compared to the results obtained with exosomes purified from stem cells [12] or plant nanovesicles [13]. Among the soluble molecules that derive from colostrum we find cytokines and growth factors, which are involved in immunomodulation [14]; in fact, as demonstrated with a quantitative proteomics analysis, exosomes from bovine colostrum are significantly enriched with proteins that can potentially regulate the immune response and growth [14]. Also, wound healing, a dynamic and complex process involving a sequence of coordinated cellular events through which the organism attempts to restore the structural integrity and complete or partial functionality of the tissue, can be improved using products based on exosomes purified from bovine colostrum [8]. The healing of skin wounds begins at the very moment of the wound and involves cells present at the site of the lesion, cells migrated from different areas of the organism (such as keratinocytes, endothelial cells, fibroblasts, inflammatory cells) and inflammatory mediators (such as cytokines, chemokines, growth factors). Many research agrees that interleukin-6 (IL-6) [15] and transforming factor beta-1 (TGF-β1) [16] play key roles in these processes. Is therefore clear that in order to promote the regeneration and healing processes of the skin it is useful to stimulate cell vitality, promote healing, stimulate the proliferation and migration of fibroblasts, endothelial cells and keratinocytes, reduce the inflammatory state and oxidative damage [17]. Similar processes are necessary for tissue rejuvenation and skin tightening [17].

The aim of this study is to compare the *in vitro* effects of some commercial products that use new technologies containing plant-derived nanovesicles or bovine colostrum-derived exosomes for facial repair, regeneration and rejuvenation.

## 2. Materials and Methods

For our research we purchased commercial products that utilize new technologies containing among the ingredients extracellular vesicles. In particular, one product contains plant extracellular vesicles extract from *Rosa damascena* callus (indicated with Rosa-NVs). The second product contains plant extracellular vesicles extract from *Centella asiatica* callus (indicated with Centella-NVs). The third product contains plant extracellular vesicles extract from *Euphorbia supina* stem (indicated with Euphorbia-NVs). The fourth compared product contains exosomes purified from bovine colostrum passively loaded with growth factors and cytokines purified from bovine colostrum (indicated with Colostrum-Exo). This latest technology is called AMPLEX plus technology.

### 2.1 Propagation and maintenance of cells

Normal adult primary human epidermal keratinocytes (HEKa) (Invitrogen) were maintained in a humidified incubator with 5% CO_2_ at 37°C in Epilife Medium containing Epilife defined growth supplement according to the manufacturer’s instructions and split every 2–3 days depending on cell confluence.

Normal adult human dermal fibroblasts (HDFa) (ThermoFisher Scientific) were cultured in Dulbecco’s Modified Eagle Medium (DMEM) supplemented with 10% FBS, streptomycin (0.3 mg mL^-1^) and penicillin (50 IU mL^-1^), and GlutaMAX (1 mM) using 75 cm^2^ polystyrene culture flasks. Cells were maintained in a humidified environment (37 °C and 5% CO_2_) and split every 2–3 days depending on cell confluence.

### 2.2 Analysis of cell proliferation/metabolic status

Keratinocytes and fibroblasts were harvested by using a trypsin-EDTA solution, counted with a hemocytometer and plated in 96-well plates (5×10^5^ keratinocytes/well; 4×10^4^ fibroblasts/well). The following day, cells were treated with four products at three different concentrations (0.5%, 1% and 2%) and incubated for 24 in a humidified environment (37 °C and 5% CO_2_). At the end of the treatment, MTT solution (1 mg/mL) was added to each well and cells were incubated for 2 hours in a humidified environment (37 °C and 5% CO_2_). Epoch Microplate Spectrophotometer (Bio Tec) was used to read the absorbance at 569 nm. Values were normalized with respect to control untreated cells and were expressed as the percent variation of cell proliferation/metabolic activity.

### 2.3 Scratch-wound assay

Human non immortalized fibroblasts were plated into six-well plates at the density of 2.5 x 10^6^ cells/well allowed to incubate until confluence. Once the confluence was reached, cells were scraped by using a 1 mL sterile pipette tip in order to form a wound. Cells were washed by using PBS to remove detached cells before adding the medium, in absence or presence of tested products used at 2%. Untreated cells were considered as control. Images were captured at 0, 1 and 24 hours after scratching.

### 2.3 Quantification and characterization of plant-derived nanovesicles and colostrum-derived exosomes

The characterization of plant-derived nanovesicles and colostrum-derived exosomes was performed with a light scattering according to [18]. Measurements were performed with a homemade apparatus using a quartz scattering cell, confocal collecting optics, a Hamamatzu photomultiplier mounted on a rotating arm, a BI-9100AT hardware correlator (Brookhaven Instruments Corporation) and illuminating the sample with a 660 nm laser. The power ranged between 5 and 15mW. Low power intensity was used to avoid convective motions due to local heating.

### 2.5 Statistical analysis

The statistical analysis was carried out by using Graphpad Prism software (version 8.0) (Graphpad software, San Diego, CA, USA). Two-way analysis of variance (ANOVA), followed by Tukey’s post hoc test, was used for multiple comparisons. The statistical significance was set at p-values < 0.01. Data were reported as the mean ± SD of at least 2 independent experiments.

## 3. Results and Discussion

The size distribution of extracellular vesicles was determined using the Dynamic Light Scattering technique. In this measurement technique, a monochromatic radiation with wavelength λ and wave vector k_l_, generally obtained via a laser, is passed through a polarizing filter and then made to impact on a sample of the colloidal dispersion of nanoparticles. When light radiation impacts a particle, scattering occurs where the light is scattered in all directions with a wave vector *k*_*f*_ which depends on the shape, dimensions and optical properties of the particle. Since the suspension is made up of a very large number *N* of particles, the intensity of the total scattering radiation that reaches the detector and is measured by it will be the sum (due to interference) of the scattering contributions of the individual particles. If we consider a colloidal dispersion, whose particles move independently with Brownian-type motion as a result of thermal agitation, the position of the particles themselves varies randomly over time. Therefore, if the light radiation emitted by scattering is collected, after being filtered by a second polarizer, by a detector, the measured intensity will fluctuate around an average value (random walking of nanoparticles) [19, 20]. In table 1 are indicated the diameter size and the concentration of the plant-derived nanovesicles and colostrum-derived exosomes. The dynamic light scattering analysis has revealed that the diameter size of Rosa-NVs (Fig 1A), Centella-NVs (Fig 1B) and Euphorbia-NVs (Fig 1C) are 565 nm, 677 nm and 667 nm respectively and that diameter size of colostrum-derived exosomes is 113 nm (Fig 1D). Exosomes are a class of cell-derived extracellular vesicles of endosomal origin and are typically 30-150 nm in diameter. They are the smallest type of extracellular vesicles, and their role is closely linked to size. Plant nanovesicles called also exosome-like are nano-sized vesicles of 50–1000 nm diameter [13]. In table 1 are indicated the diameter size and the concentration of the plant-derived nanovesicles and colostrum-derived exosomes of tested products. The dynamic light scattering analysis has revealed that the diameter size of Rosa-NVs (Fig 1A), Centella-NVs (Fig 1B) and Euphorbia-NVs (Fig 1C) are 565 nm, 677 nm and 667 nm respectively and that diameter size of colostrum-derived exosomes is 113 nm (Fig 1D). Exosomes are a class of cell-derived extracellular vesicles of endosomal origin and are typically 30-150 nm in diameter. They are the smallest type of extracellular vesicles, and their role is closely linked to size. Plant nanovesicles called also exosome-like are nano-sized vesicles of 50–1000 nm diameter [13]. Recent data have suggested that plant exosome-like nanovesicles (PENVs) and PENV-derived artificial nanocarriers (APNVs) may have possible clinical applications, but there are many obstacles to overcome [13]. Currently extracted plant nanovesicles are highly heterogeneous, containing multiple vesicle types, and of unknown origin. originates in plant cells [13]. Another problem to overcome depends on the fact that, because PENVs and APNVs do not come from mammalian cells, they may have poor targeting ability in some targeted tissues [13].

**Table 1.** Diameter size and concentration of extracellular vesicles analyzed with DLS.

| Product | Diameter size (nm) | Concentration (particles/ml) |
| --- | --- | --- |
| Rosa-NVs | 565 | $2^6$ |
| Centella-NVs | 677 | $2^6$ |
| Euphorbia-NVs | 667 | $2^5$ |
| Colostrum-Exo | 113 | $3.8^9$ |

**Figure 1.**
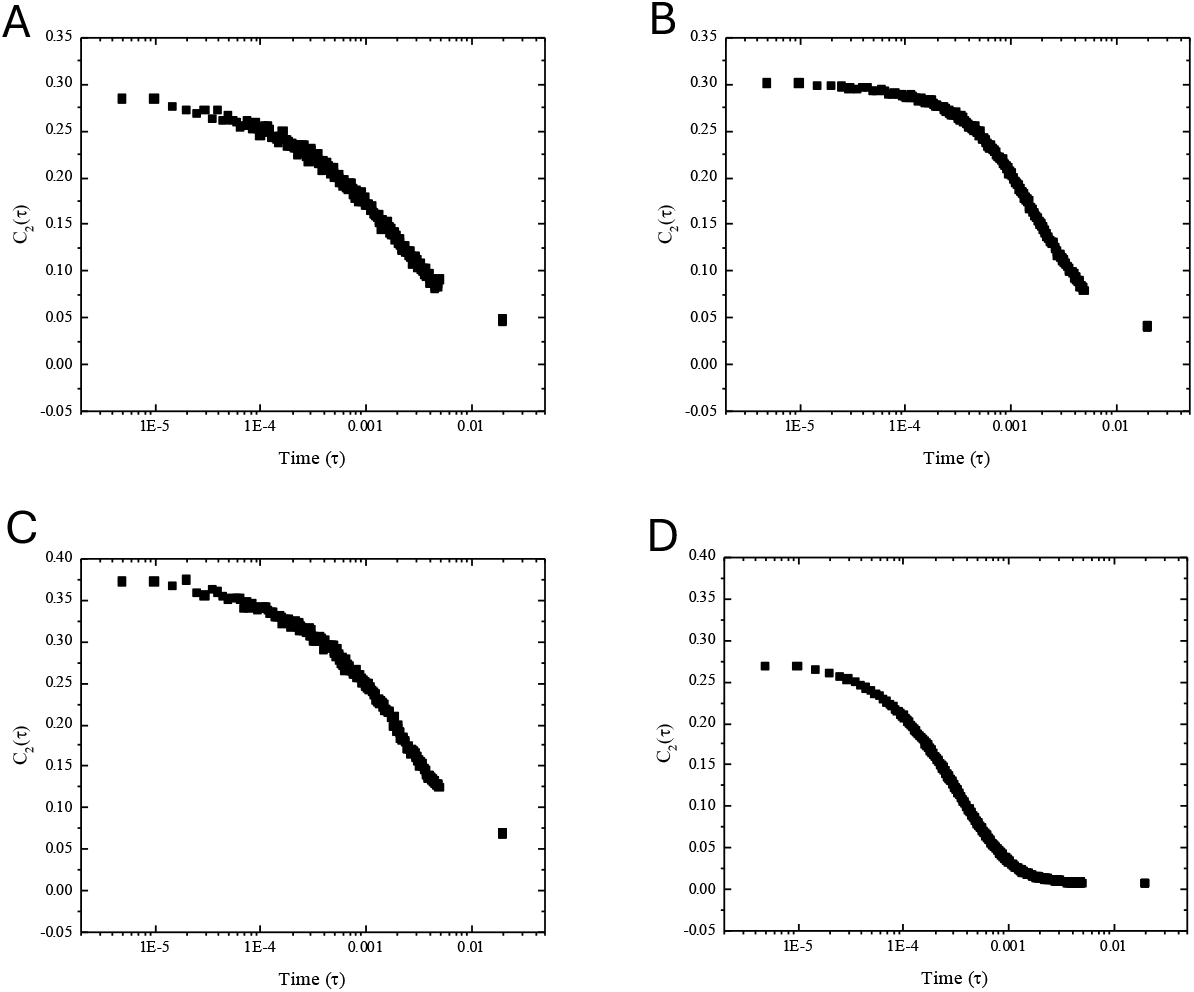
DLS analysis used to characterize the size distributions of exosomes within four products: (A) product with plant extracellular vesicles from *Rosa damascena* callus; (B) product with plant extracellular vesicles from *Centella asiatica* callus; (C) product with plant extracellular vesicles from *Euphorbia supina* stem; (D) product with exosomes from colostrum bovine loaded with growth factors and cytokines purified from bovine colostrum.

In the four products analyzed the concentration of nanovesicles was also different (Table 1), lower for plant-derived nanovesicles (2^6^ part/ml, 2^6^ part/ml and 2^5^ part/ml for Rosa-NVs, Centella-NVs and Euphorbia-NVs respectively) and higher for colostrum-derided exosomes. (3.8^9^ part/ml).

The viability cell assay was conducted for both, human keratinocytes and dermal fibroblasts, and the results are presented in Figures 2 and 3, respectively. The results showed a statistically significant increase in the level of cell viability after treatment with product containing colostrum exosomes at different doses (p<0.001) compared to cells treatment with plant-derived nanovesicles, whose results were lower than the negative control (untreated cells) at all concentrations tested.

**Figure 2.**
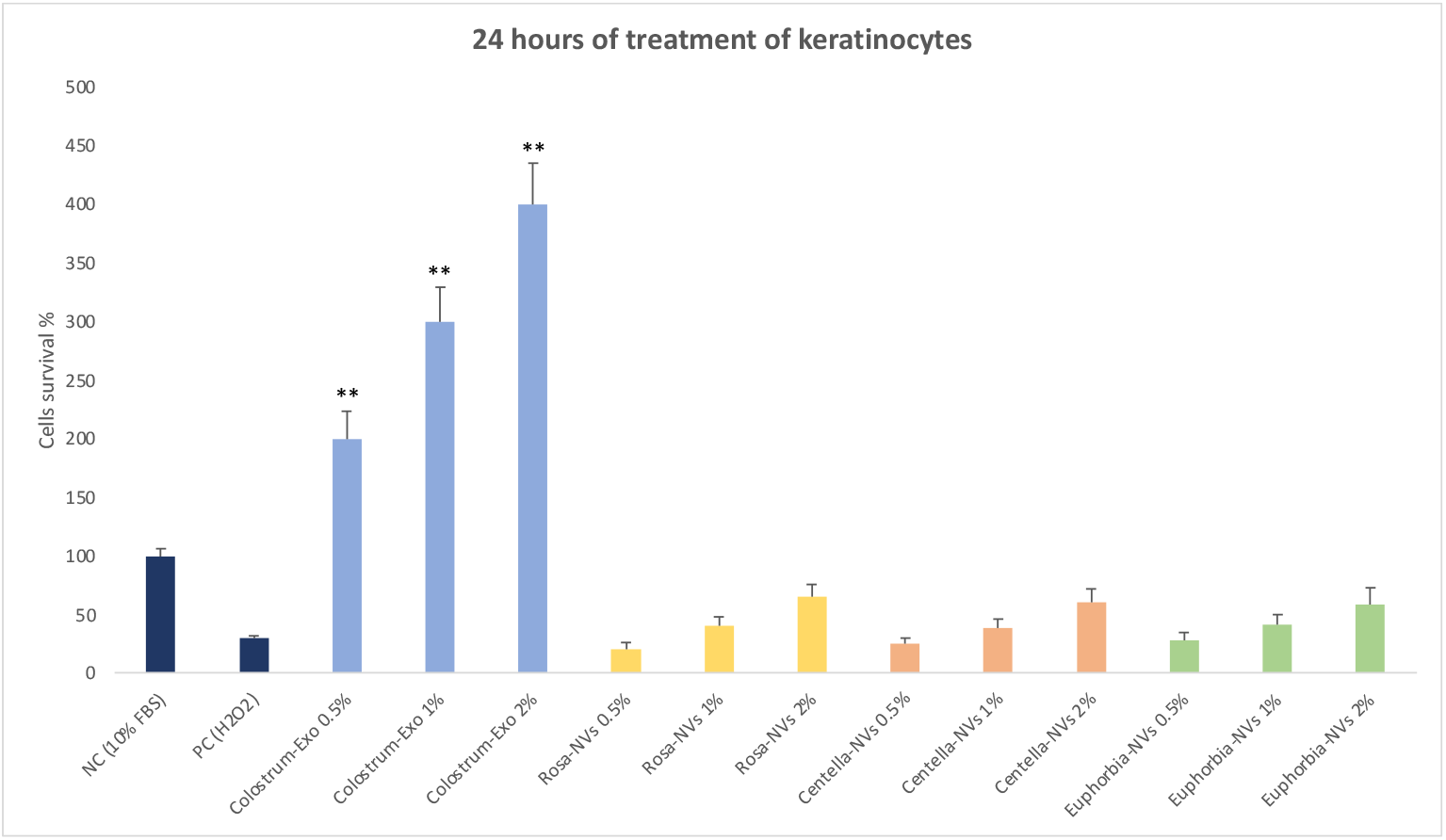
The cell survival of human keratinocytes at 24h following treatment with the four products tested to different concentrations (0.5%, 1% and 2%). NC (Negative Control, cells without treatment); PC (Positive Control, 80 µM H_2_O_2_); The asterisks denote the degree of significance between results: **p<0.001. Errors bars represent the Standard Deviation of the mean (experiment was repeated 2 times).

**Figure 3.**
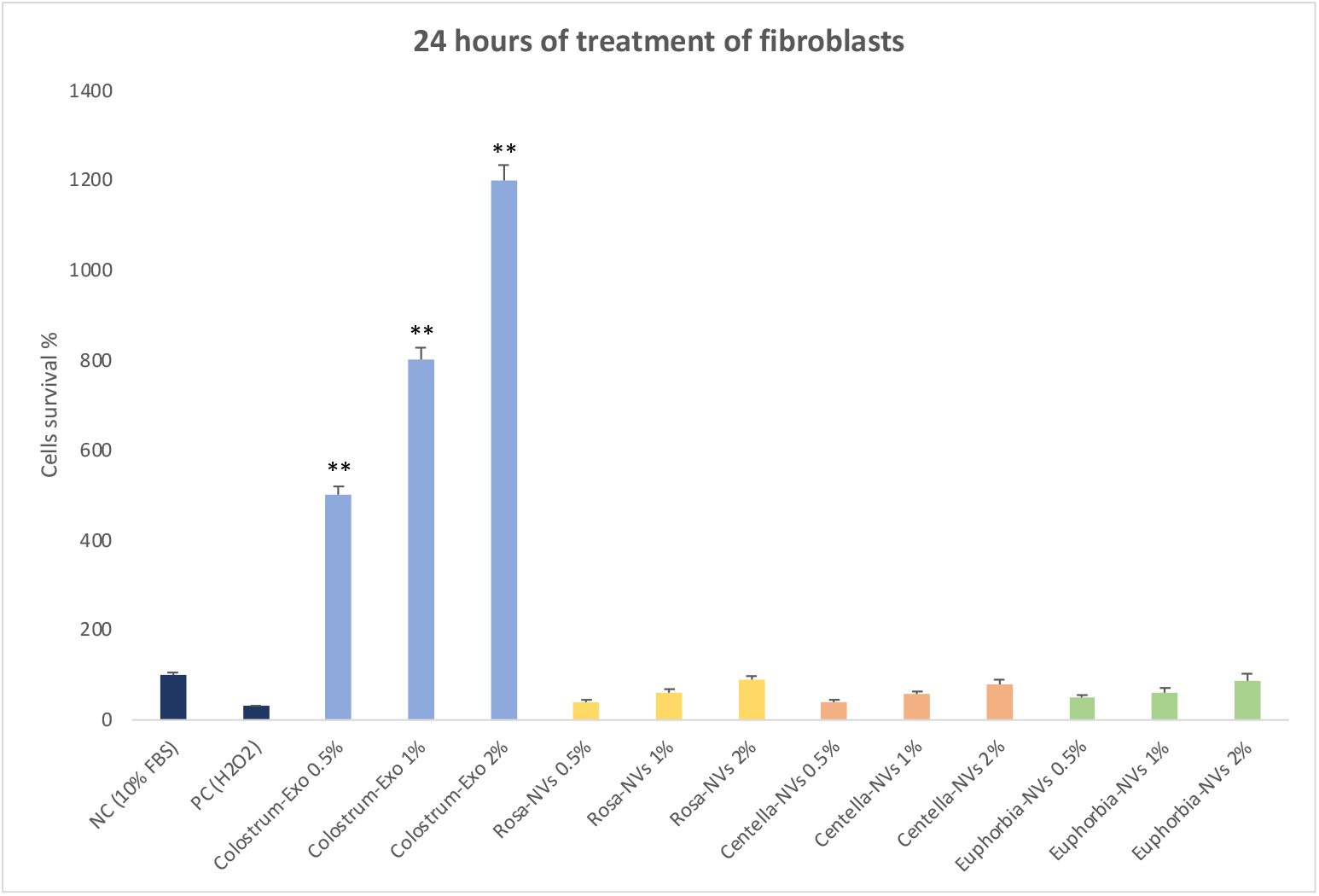
The cell survival of human fibroblasts at 24h following treatment with the four products tested to different concentrations (0.5%, 1% and 2%). NC (Negative Control, cells without treatment); PC (Positive Control, 80 µM H_2_O_2_); The asterisks denote the degree of significance between results: **p<0.001. Errors bars represent the Standard Deviation of the mean (experiment was repeated 2 times).

The experimental results obtained with the scratch test have demonstrated a significant increase in the level of cell proliferation following a 1 hour and 24 hour treatment with product containing colostrum-derived exosomes loaded with colostrum biomolecules (growth factors and cytokines) at 2% concentrations. Figure 4, in fact, show a statistically significant increase in the level of cell viability after treatment with product containing colostrum exosomes (p<0.001) compared to cells treatment with plant-derived nanovesicles and negative control (untreated cells).

**Figure 3.**
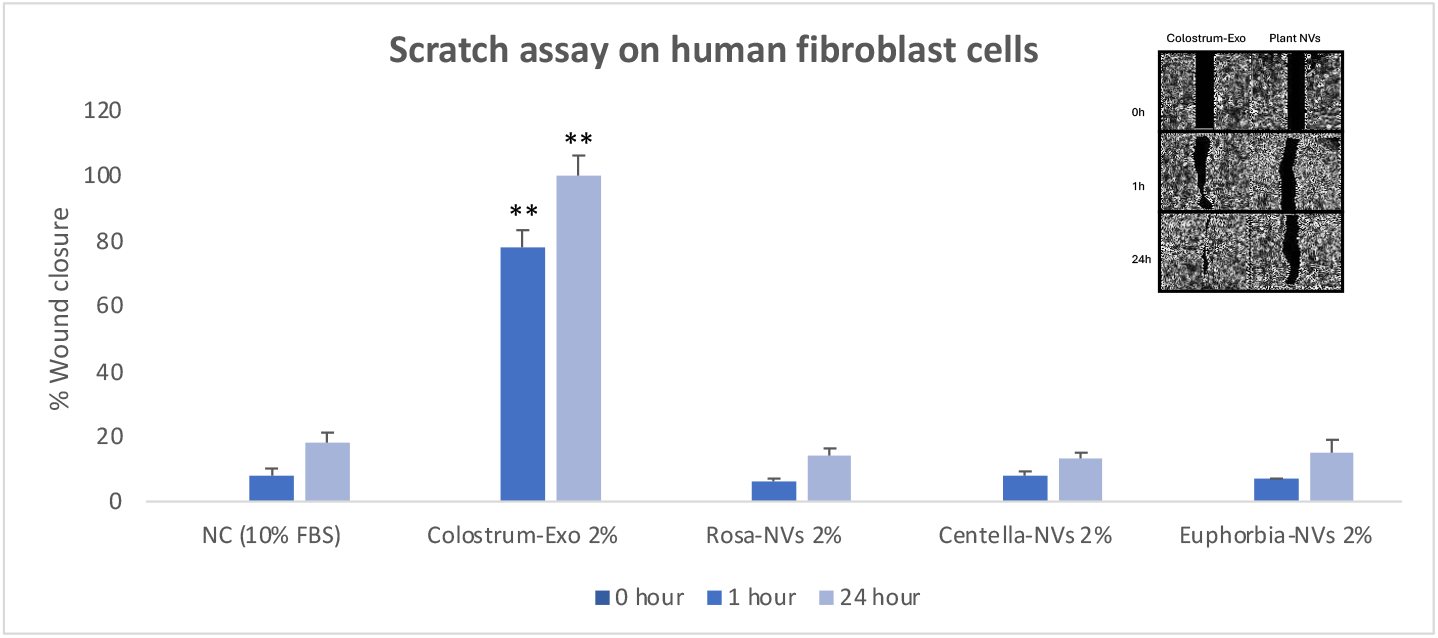
Scratch assay on human fibroblasts treated with the four products to 2% concentration at 0, 1 and 24 hours’ time points. Untreated cells were considered as a control (NC). The asterisks denote the degree of significance between results: **p<0.001. Errors bars represent the Standard Deviation of the mean (experiment was repeated 3 times). The box shows the images comparing the treatment with colostrum-derived exosomes and plant-derived nanovesicles (the results were comparable for the three products containing plant NVs).

Wound healing, as well as tissue rejuvenation and skin tightening, are a complex processes that involves multiple stages, hemostasis, inflammation, proliferation, and tissue regeneration being the most critical steps [21]. The focus of this study was to examine the *in vitro* effects of some commercial products that use new technologies containing plant derived nanovesicles or exosomes for facial repair, regeneration and rejuvenation. The results obtained have demonstrated that the product containing colostrum-derived exosomes passively loaded with growth factors and cytokines from colostrum (AMPLEX plus technology) promote cell vitality and wound healing, determining a statistically significant increase in the level of cell viability after treatment at different doses (0.5%, 1% and 2%) compared to cells treatment with plant-derived nanovesicles. Also the results obtained with the scratch test have demonstrated a significant increase in the level of cell proliferation already after a 1 hour treatment (78%) with product containing colostrum-derived exosomes loaded with colostrum biomolecules (growth factors and cytokines) at 2% concentrations. Our results are in according to other studies published in which the effectiveness of plant-derived nanovesicles was analyzed on facial repair, regeneration and rejuvenation. The results obtained in these studies have showed a low efficacy of plant nanovesicles, almost always comparable to untreated cells or even lower [13, 22-26]. Some Authors, for example, hypothesized that *Rose damascena* stem cell in culture can release exosome-like particles that may have biological function in cells relevant to skin, such as hair papilla cells, fibroblasts and melanocytes [23]. The Authors have found that the Rose Stem Cell cultured supernatant reduce viability in human dermal papilla cells, demonstrating its cytotoxic effect. However, the *Rose damascena* stem cell nanovesicles did not show any detrimental effect on viability of Human Dermal Papilla cells, but they have not even demonstrated a positive effect. Furthermore, the Authors declare that the *Rose damascena* stem cell nanovesicles also stimulate skin fibroblast proliferation and collagen production, as well as *in vitro* wound healing, but their results are just higher than those of untreated cells and after 24 hours there was a closure of just 27.5% [23], much lower than obtained in our experiments with bovine colostrum exosomes-derived after only one hour of treatment, equal to 78%. Other Authors to confirm the cellular toxicity of *Centella asiatica* extract and *Centella asiatica* exosomes, have performed the MTT assay with a keratinocyte cell line (KEKa cells) [24]. Centella extract and Centella exosomes were nontoxic to the cells at all concentrations and induced cell proliferation compared with the control. Moreover, Centella extract induced higher proliferation than Centella exosomes; in particular, Centella exosomes has a maximum percentage of cell viability equal to 123.7 ± 4.8 % with a tested concentration of 1x10^5^ particles/ml. In literature there is no research on the effects of exosomes purified from *Euphorbia asiatica*, but some authors have studied the antioxidant and skin-whitening effects of a 70% ethanol extract of Euphorbia extract. Authors have showed that Euphorbia extract significantly reduced tyrosinase activity and melanin content in a dose-dependent manner and MTT test it showed that the Euphorbia extract was not toxic as the percentage cell viability was the same to that of untreated cells [25]. Exosomes from colostrum bovine instead have an excellent structural and have a great potential as natural therapeutic agents to repair UV-irradiated skin aging and damage [27]. Authors showed that treatment with colostrum exosomes prevented the UV-induced generation of intracellular reactive oxygen species in epidermal keratinocytes. In UV-stimulated melanocytes, have reduce significantly the production of the protective skin-darkening pigment melanin; in the human dermal fibroblasts treated with colostrum exosomes, the expression of matrix metalloproteinases was suppressed, whereas increased cell proliferation was accompanied by enhanced production of collagen, a major extracellular matrix component of skin [27]. Other Authors suggested its potential antiaging properties. These effects are attributed to its capacity for moisturization and wrinkle reduction, which correlate with improvements in the expression levels of FLG, CD44 in keratinocytes, and HAS2 in fibroblasts [28].

## 4. Conclusion

Extracellular vesicles, including exosomes, are produced via the endosomal pathway and released in the extracellular space upon fusion with the plasma membrane [29]. Several studies show that these extracellular vesicles play a key role in cell-to-cell communication, transporting bioactive molecules [30-32]. Exosomes were reported to regulate also all phases of skin repair and rejuvenation [33, 36]. In particular, exosomes from colostrum they appear to have a high potential for the applications in regenerative medicine [27, 28], as also demonstrated by our research. Of all the products tested, in fact, only product containing exosomes purified from colostrum passively loaded with growth factors and cytokines derived purified from colostrum (AMPLEX plus technology) gave very encouraging results in terms of effects on proliferation, cell viability and wound repair. The other products tested gave results comparable to the untreated samples, or even lower.

## Author Contributions

Conceptualization, M.V.B.; methodology, M.V.B.; software, G.F. and M.Z.; formal analysis, G.F. and M.Z.; investigation, G.F.; data curation, A.P., G.F. and M.Z.; writing original draft preparation, M.V.B.; Writing – Review & Editing A.P.; supervision, M.V.B. All authors have read and agreed to the published version of the manuscript.

## Institutional Review Board Statement

This study was performed in line with the principles of the Declaration of Helsinki and does not require approval from the Ethics Committee of University of Catania.

## Acknowledgments

G.F. thanks the Ph.D. program FSE Notice 1/2021.

